# Glutamate delta-1 receptors regulate a novel tonic excitatory conductance in the mouse bed nucleus of the stria terminalis and influence neuronal function

**DOI:** 10.1101/2025.04.15.649028

**Authors:** Sara Y Conley, Sarah E Sizer, Madigan L Bedard, Zoé A McElligott

## Abstract

The plasticity of ionotropic receptors (mainly AMPARs and NMDARs) within the glutamatergic system has long been investigated as a mechanism for physiological and pathological adaptive learning. The tetrameric delta glutamate receptors (δGluRs, GluD1 and GluD2) are homologs of the AMPARs and NMDARs, however they are insensitive to glutamate. These proteins, especially GluD1, have been implicated in multiple psychiatric conditions and play a functional role in synapses assembly and stability, but recent evidence suggests they also may supply a tonic excitatory conductance, are sensitive to poly-amine blockade, and regulate synaptic plasticity. Here we use whole-cell patch clamp electrophysiology to functionally characterize these receptors in the dorsolateral bed nucleus of the stria terminalis (dlBNST) and investigate their modulation of both synaptic transmission and cell excitability. Our results suggest that δGluRs, and in particular GluD1, carry a tonic conductance, modulate excitatory synapses, and regulate cell excitability in the dlBNST. These results imply that GluD1 functions to regulate the flow of information through the BNST and may play a role in affect, stress, and substance use disorders.

## Introduction

Glutamate, the principal excitatory neurotransmitter in the central nervous system, mediates a variety of physiological processes through a diverse array of receptors. Among these, ionotropic glutamate receptors (iGluRs) play critical roles in synaptic transmission, plasticity, and neurodevelopment. While much attention has been devoted to the well-characterized NMDA, AMPA, and kainate receptors, a lesser-studied subtype, the delta glutamate receptors (δGluRs), have emerged as an intriguing component of the “glutamatergic system”.

δGluRs, GluD1 and GluD2, tetrameric homomers so categorized due to their sequence homology with other ionotropic glutamate receptors, are unique in that they do not respond to glutamate, and their endogenous ligand remains unclear (Yuzaki and Aricescu 2017; Burada et al. 2022), although several candidate molecules have been proposed including D-serine, glycine, and GABA (Naur et al. 2007; Carrillo et al. 2021; Piot et al. 2023). Instead, δGluRs appear to mediate synapse stabilization by complexing with synaptic structural proteins cerebellin and neurexin, and facilitate synaptic plasticity (Dai et al. 2021). The most well-studied example of this is in the cerebellum, where GluD2 is primarily expressed; GluD2 physically interacts with the Gq-coupled mGluR1 to induce the internalization of GluA1-containing calcium-permeable AMPA receptors (CP-AMPARs), resulting in long-term depression (LTD) of parallel fiber-Purkinje cell synapses (Ito 2002; Ichikawa et al. 2016). Additionally, mGluR1 has been shown to gate the opening of GluD2, causing a slow excitatory postsynaptic current (EPSC) (Ady et al. 2014). GluD1, which is more ubiquitously expressed throughout the brain (Nakamoto et al. 2020b), has been shown to function similarly as an effector for Gq-coupled receptors. GluD1 knockouts (KOs) display aberrant mGluR5-dependent LTD in the hippocampus (Suryavanshi et al. 2016), while in the dorsal raphe, GluD1 mediates the α1-adrenergic receptor (α1-AR) dependent slow EPSC (Gantz et al. 2020). Recent studies have also demonstrated evidence for constitutively active GluD1 in the dorsal raphe, conferring a tonic excitatory current that is blocked by the polyamine 1-naphthylacetyl spermine trihydrochloride (NASPM), a drug canonically used to block CP-AMPARs (Gantz et al. 2020; Copeland et al. 2023). Despite their emerging importance, the precise mechanisms of δGluR activation, signaling pathways, their contribution to synaptic plasticity, and how these may vary across brain regions remain poorly understood. The bed nucleus of the stria terminalis (BNST) is a forebrain nucleus reciprocally connected to many important stress and reward-related structures, positioning it as a critical nexus through which the negative affective symptoms common to many psychiatric- and substance use-disorders are mediated (McElligott and Winder 2009; Gungor and Paré 2016). Studies have shown alterations in BNST plasticity and function following stress and drug and alcohol exposure (Ch’ng et al. 2018). Moreover, the BNST neurons express multiple Gq-coupled LTDs of excitatory transmission that are induced and maintained by different mechanisms (McElligott et al. 2010). Here signaling at α^1^-adrenergic receptors (ARs) induces an LTD maintained in part by internalization of NASPM-sensitive CP-AMPARs, which contribute substantially to glutamatergic synapses in naive male mice. While previous studies have shown that the dorsolateral (dl) BNST highly expresses GluD1(Konno et al. 2014; Nakamoto et al. 2020b), the functional role of GluD1 in these neurons has not previously been assessed. In probing dlBNST CP-AMPARs we serendipitously discovered that NASPM also shifted the holding current of the cells. In this study, we followed up on this discovery using whole-cell patch clamp electrophysiology in naive male C57BL6/J, and GluD1 knockout (KO) and wild-type (WT) littermates. We show that the dlBNST expresses a NASPM-sensitive, GluD1-mediated tonic excitatory conductance which influences cell excitability and AMPAR currents. These experiments establish the role of tonic excitatory conductance within the BNST and set the stage for future studies probing this phenomenon in modes of stress and substance use disorders.

## Methods

### Mice

All mice used in these experiments were males. C57BL/6J mice aged 8-12 weeks were purchased from The Jackson Laboratory (Strain #:000664). GluD1 KOs and WT littermates are derived from GluD1 breeders originally generated by Dr. Jian Zuo (Gao et al. 2007). All mice were at least 8 weeks old prior to any procedures. Mice were group housed (2-5 per cage), maintained on a normal 12-hour light-dark cycle (lights on 7:30 AM/lights off 7:30 PM) and had access to food and water *ad libitum*. All procedures described were approved by the University of North Carolina at Chapel Hill Institutional Animal Care and Use Committee (IACUC).

### Brain Slice Preparation

After retrieval from the UNC Animal Facility, mice were allowed to rest for a minimum of 30 minutes in sound and light-attenuating chambers. Mice were then anesthetized with isoflurane and decapitated. Brains were extracted, hemisected, and placed into oxygenated (95% vol/vol O2, 5% vol/vol CO_2_), high-sucrose, low-Na^+^ ACSF maintained at 1–4 °C (sucrose ACSF: 194 mM sucrose, 20 mM NaCl, 4.4 mM KCl, 2 mM CaCl2, 1 mM MgCl2, 1.2 mM NaH2PO4, 10 mM glucose, 26 mM NaHCO3), where 250 μm thick coronal slices were collected using a Leica VT1000S vibratome (Leica Microsystems). Following slicing, brains were allowed to equilibrate in standard aCSF (in mM: 124 NaCl, 4.4 KCl, 2 CaCl_2_, 1.2 MgSO_4_, 1 NaH_2_PO_4_, 10 glucose, 26 NaHCO_3_, 32°C) for at least 30 min before recording.

### Whole Cell Electrophysiology

Slices were then transferred to the recording chamber and allowed to equilibrate in oxygenated aCSF containing 25 µM picrotoxin (PTX), maintained at 30°C and perfused at 2 ml/min, for an additional 30 min before recordings were initiated to ensure full saturation of PTX. Patch electrodes (3– 6 MΩ) were pulled using a Flaming/Brown micropipette puller (P-97 Sutter Instruments) and filled with either Cs-gluconate or K-gluconate internal solution. Voltage-clamp recordings were conducted using Cs-gluconate (in mM: 135 Cs-gluconate, 5 NaCl, 10 HEPES, 0.6 EGTA, 4 ATP, 0.4 GTP). For experiments examining sEPSCs, cells were voltage-clamped at −80 mV to increase driving force of ions through the glutamate receptors. Recordings examining cell excitability were performed in current-clamp using K-gluconate intracellular recording solution (in mm: 135 K-gluconate, 5 NaCl, 2 MgCl_2_, 10 HEPES, 0.6 EGTA, 4 Na_2_ATP, 0.4 Na_2_GTP). All data were acquired using Clampex 11 (Molecular Devices). Access resistance (AR) was monitored continuously, and recordings for which AR changed >20% were excluded from analysis. All data were analyzed using either Clampfit 11 (Molecular Devices) or Easy Electrophysiology (Easy Electrophysiology Ltd). Holding current/resting membrane potential values were calculated in Clampfit 11 using the mean of a Gaussian fit to an all-points histogram of the trace in the final minute of baseline and of drug application (Glykys and Mody 2007; Zhu et al. 2023).

### Statistics, Rigor, and Reproducibility

All statistics were calculated using GraphPad Prism v10. Data are reported as groups means and/or individual data points ± SEM unless otherwise noted in the figure legend. Comparisons were made using one-sample t-tests, Student’s t-test, or 2-way ANOVAs as appropriate. Correlations were determined by calculating the Pearson Correlation Coefficient. Significance was defined as p ≤ 0.05. Symbols for statistical significance follow GraphPad p-value style (p > 0.05 ns, p ≤ 0.05 *, p ≤ 0.01 **, p ≤ 0.001 ***, p ≤ 0.0001 ****).

## Results

### NASPM decreases tonic and synaptic excitatory signaling in naïve C57BL6/J males

Figure 1 demonstrates the existence of a NASPM-sensitive tonic excitatory current in the dlBNST of naïve male C57BL6/J mice. Application of 100 µM NASPM caused a significant upward shift in holding current compared to baseline (Fig. 1b), with an average magnitude (Fig. 1c) of 20.03 pA (t_(10)_=6.144, p=0.0001). While we observed a positive trend between baseline holding current and tonic current magnitude (Fig. 1d), this correlation did not reach statistical significance (R_2_=0.2624, p=0.1072). We next evaluated the effect of NASPM on spontaneous excitatory synaptic transmission in the dlBNST of naïve C57BL6/J mice (Fig. 2a-e). NASPM significantly decreased sEPSC frequency (t_(10)_=2.750, p=0.0205), but not amplitude (t_(10)_=1.889, p=0.0883). We calculated a normalized excitatory drive using the formula 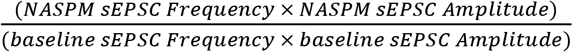, and found this to be significantly reduced as a result of NASPM as well (t_(10)_=3.610, p=0.0048). No significant changes were observed in sEPSC kinetics (Supplemental Fig. 1).

**Fig. 1.**
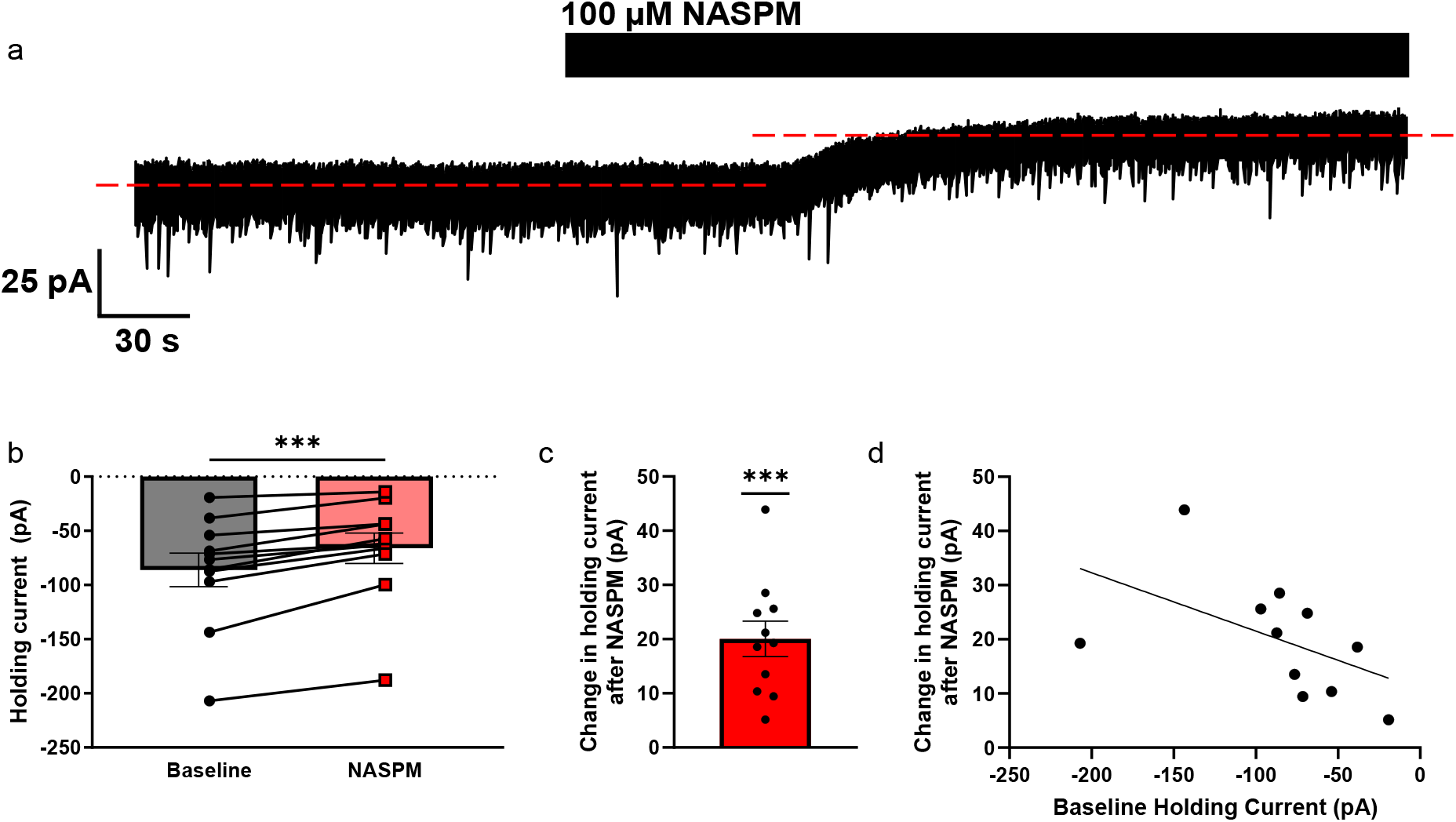
Application of 100 µM NASPM blocks a tonic excitatory conductance in the dlBNST of naïve C57BL6/J male mice. **a**. Representative voltage-clamp trace of NASPM-induced upward shift in holding current. **b**. Raw holding current values before and after NASPM application. Asterisks indicate significant difference in paired t-test **c**. Magnitude of NASPM-sensitive tonic current. Asterisks indicate significant difference in one-sample t-test compared to a theoretical mean of 0. **d**. Correlation of holding current at baseline and tonic current magnitude. Total n=11 cells from 8 mice.

**Fig. 2.**
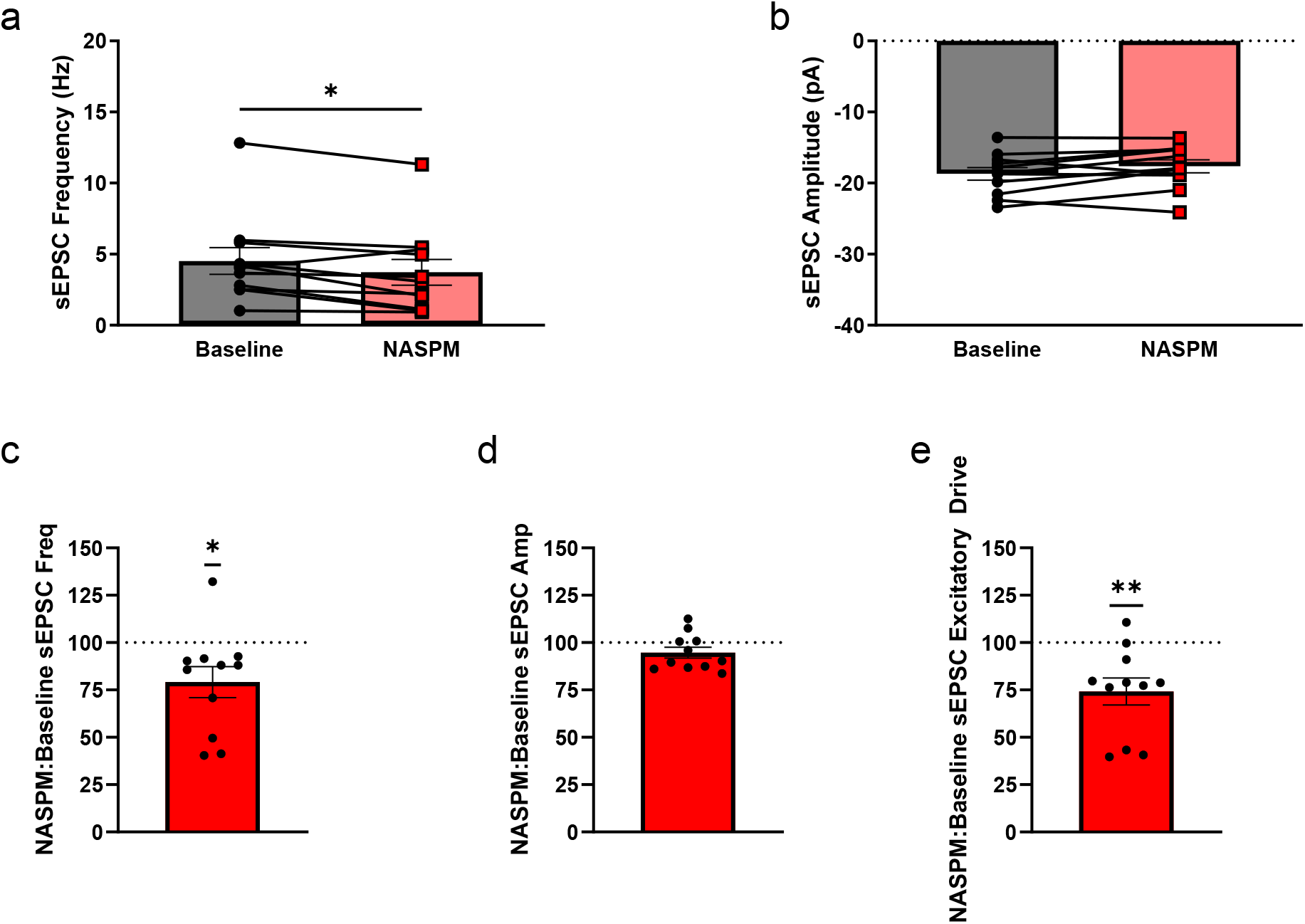
Application of 100 µM NASPM decreases synaptic excitatory signaling in the dlBNST of naïve C57BL6/J male mice. NASPM significantly decreases the frequency (**a**,**c**) but not amplitude (**b**,**d**) of sEPSCs, leading to an overall decrease in excitatory drive (**e**). Total n=11 cells from 8 mice. Asterisks indicate significant difference in paired t-tests (**a**,**b**) or one-sample t-tests (**c-e**) with a theoretical mean of 100.

### Testing NASPM effect on resting membrane potential and firing properties in naïve C57BL6/J males

A reduction in the holding current of a cell implies hyperpolarization in the resting membrane potential (RMP) as a result of NASPM application, which may impact cell excitability. We conducted current-clamp recordings next to directly test whether this was the case. Indeed, we found that NASPM significantly hyperpolarized dlBNST cell RMP (Fig. 3b) by an average −3.96 mV (Fig. 3c) (t_(12)_=3.913, p=0.0021). Further, this increased the rheobase of these cells (Fig. 3d) (t_(12)_=2.253, p=0.0438), and two-way ANOVA revealed a main effect of NASPM attenuation of the number of action potentials fired in response to 20 pA injections (from 0-200 pA) (Fig. 3e, F_(1,11)_=5.205, p=0.0434). Together, these results suggest that the NASPM-sensitive tonic excitatory conductance we observed in our voltage-clamp recordings provides meaningful physiological contribution to the intrinsic excitability of dlBNST neurons.

**Fig. 3.**
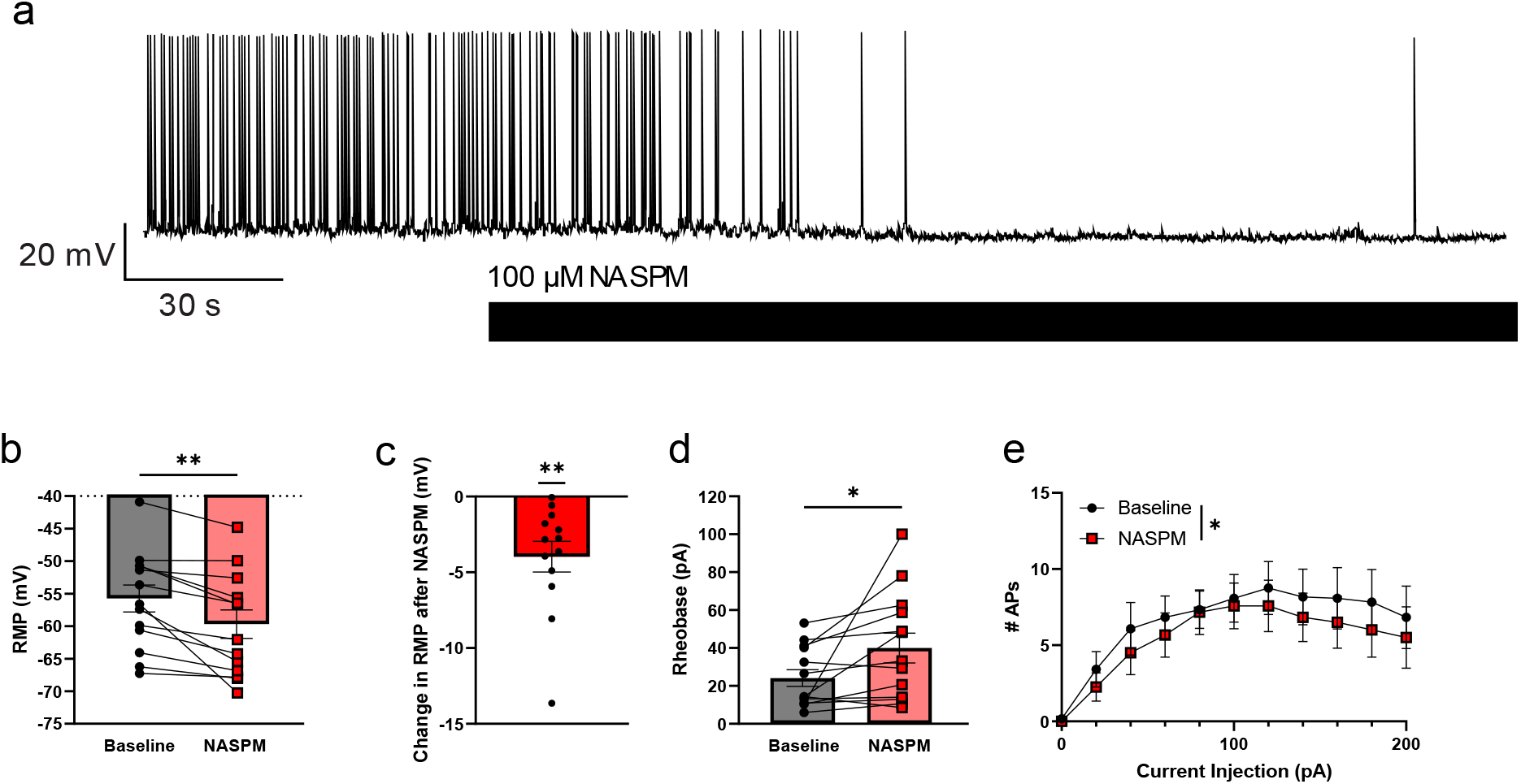
Application of NASPM hyperpolarizes the resting membrane potential and decreases excitability of dlBNST neurons in C57BL6/J male mice. **a**. Representative trace of RMP hyperpolarization after NASPM application. **b**. Raw change in resting membrane potential before and after NASPM. **c**. Average magnitude of NASPM-induced hyperpolarization. **d**. NASPM causes an increase in the rheobase at RMP. **e**. No significant changes in the number of action potentials fired by dlBNST neurons in response to increasing 20 pA current steps before and after drug application. Total n=13 cells from 6 mice. Asterisks indicate significant differences in paired t-tests (**b**,**d**), one-sample t-tests (**c**) with a theoretical mean of 0, or two-way repeated measures ANOVA with Greenhouse-Geisser correction (**e**).

### GluD1 knockout eliminates tonic excitatory conductance in the dlBNST of male mice

Since this NASPM-sensitive conductance showed similar characteristics of the GluD1-dependent tonic current that had been characterized in the dorsal raphe, we next tested to see whether knockout of GluD1 would attenuate the effects of NASPM in the dlBNST. Recording in whole-cell voltage clamp we again recorded baseline currents, and then bath applied 100 µM NASPM to the slice. Two-way RM ANOVA showed a main effect of drug (F_(1,29)_=27.22, p<0.0001), genotype (F_(1,29)_=6.100, p=0.0196), and a drug x genotype interaction (F_(1,29)_=9.606, p=0.0043), indicating that GluD1 KO mice had smaller holding currents at baseline compared to WT littermates, and NASPM caused a significant reduction in the holding current in WT mice but failed to elicit a change in this holding current in the KO animals (Fig. 4c,d). Additionally, there was a significant correlation between the baseline holding current and the magnitude of the tonic excitatory current in the WT (R_2_=0.3158, p=0.0235), but not the GluD1 KO, mice (R_2_=0.06949, p=0.3425) (Fig. 4e). When we looked at the effects of GluD1 knockout on NASPM-induced changes in synaptic transmission (Fig. 5a-e), we found that while sEPSC frequency was significantly decreased after NASPM in both genotypes (F_(1,29)_=17.37, p=0.0003), there was a greater reduction in the GluD1 KOs compared to the WTs (t_(29)_=2.549, p=0.0129). However, NASPM significantly alter the sEPSC amplitude (Fig. 5b) or kinetics (Supplemental Fig. 2) in either genotype, consistent with our observations in C57BL6/J animals. Thus, NASPM application decreases excitatory transmission in both genotypes, which was exacerbated in the GluD1 KOs (t_(29)_=2.482, p=0.0191).

**Fig. 4.**
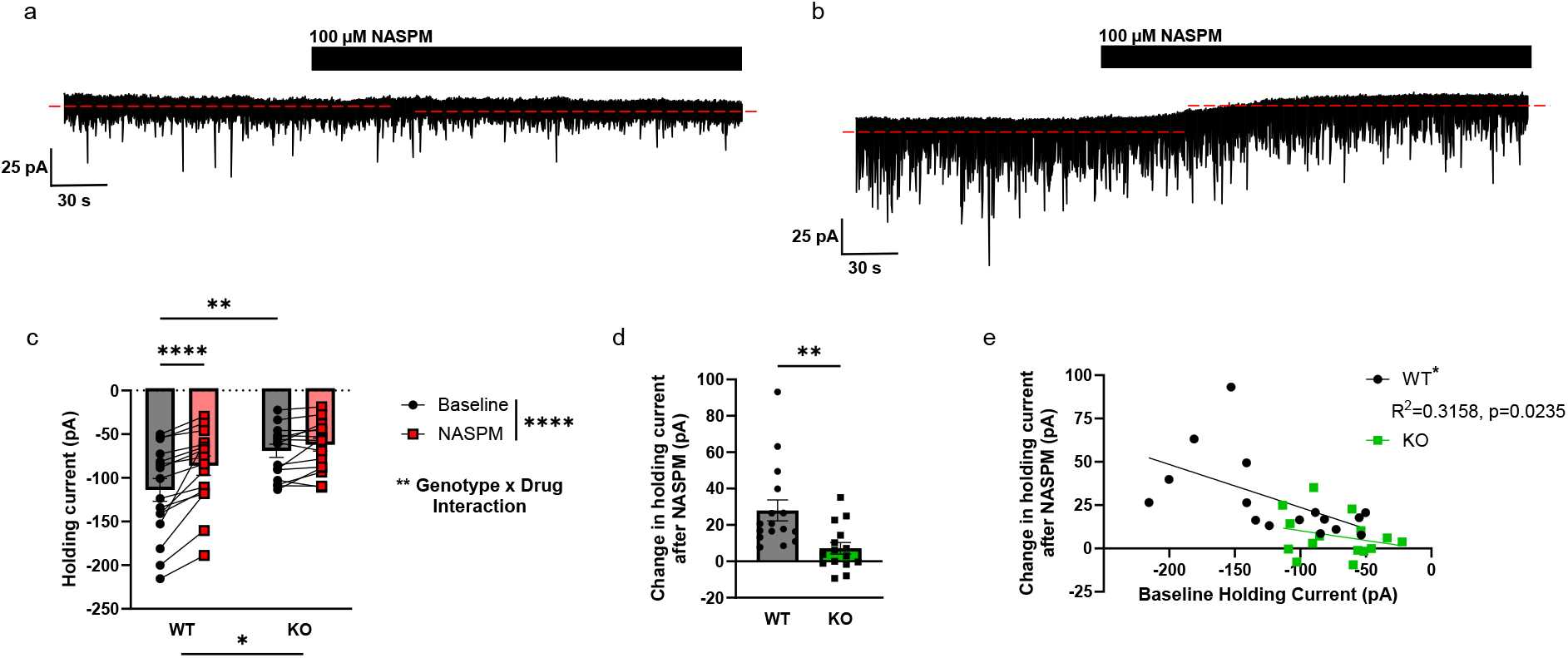
Constitutive knockout of GluD1 receptors attenuates tonic current. **a**. Representative trace of dlBNST neuron from GluD1 KO mouse. **b**. Representative trace of dlBNST neuron from WT littermate. **c**. NASPM application significantly reduces holding current in the dlBNST of WT, but not in GluD1 KO mice. **d**. Average magnitude of holding current shift from NASPM. **e**. Correlation of holding current at baseline and tonic current magnitude in WT (black, significant correlation) and GluD1 KO (green, no significant correlation). Total n = 16 cells, 9 mice for WTs and n= 15 cells, 7 mice for KOs. Asterisks indicate significant differences determined via two-way repeated measures ANOVA and post-hoc differences determined by Fisher’s LSD (**c**), unpaired t-test (**d**), or Pearson’s Correlation Coefficient (**e**).

**Fig. 5.**
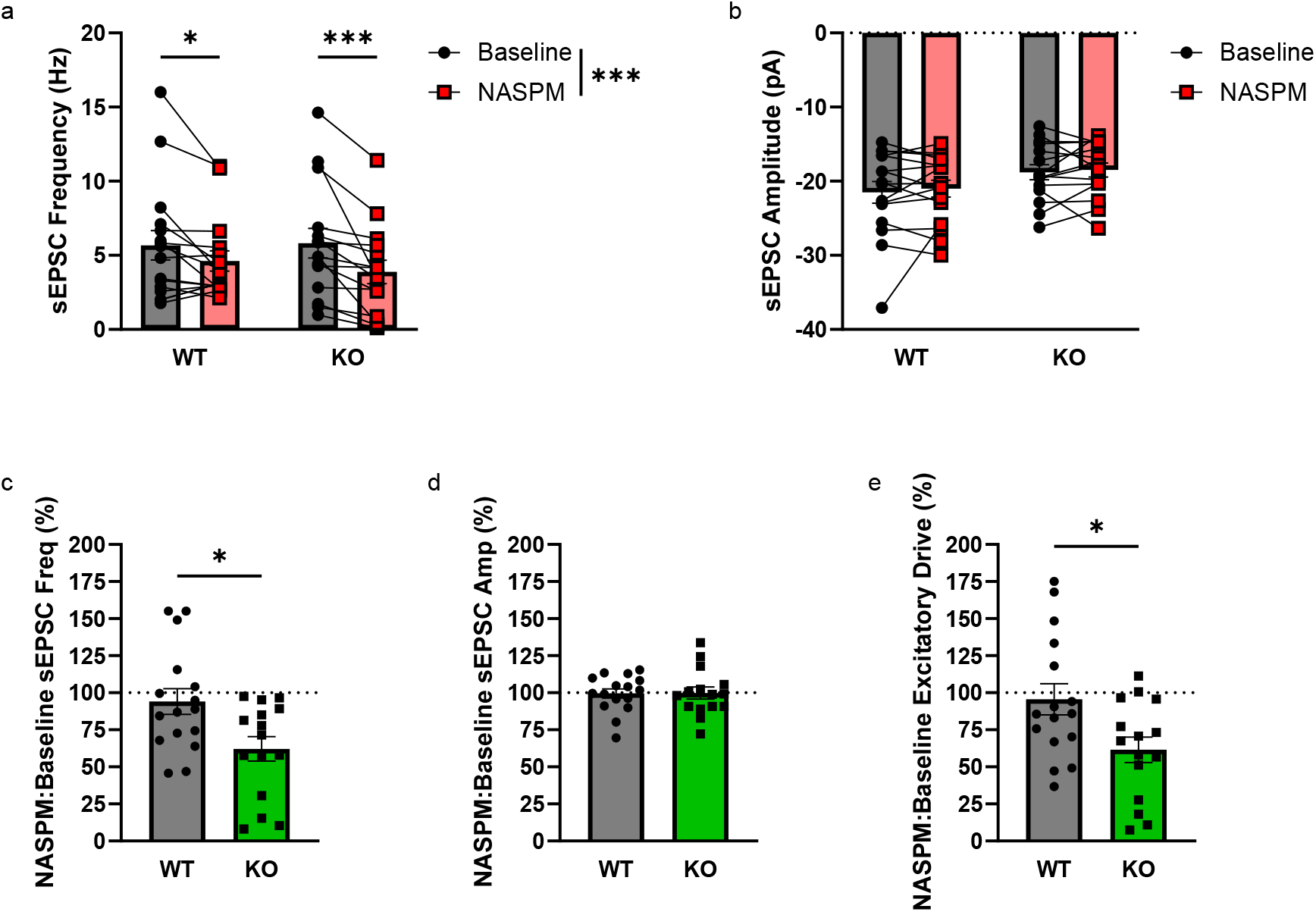
Constitutive knockout of GluD1 receptors exacerbates NASPM-mediated reductions in synaptic excitatory signaling. **a**. 100 µM NASPM significantly reduces sEPSC frequency in both genotypes. **b**. NASPM does not change sEPSC amplitude in either genotype. **c**. NASPM causes greater reduction to sEPSC frequency in GluD1 KO animals compared to WTs. **d.** No significant decrease to normalized amplitude occurs after NASPM. **e**. NASPM reduces total excitatory drive in both genotypes compared to baseline. Total n = 16 cells, 9 mice for WTs, 15 cells, 7 mice for GluD1 KOs. Asterisks are indicative of significance as determined by two-way RM ANOVA and post-hoc Fisher’s LSD (**a**,**b**) or unpaired t-tests (**c-e**).

## Discussion

The present study provides novel insights into the role of δ-glutamate receptors (δGluRs), specifically GluD1, in modulating excitatory synaptic transmission and neuronal excitability in the dlBNST. After applying NASPM, we observed significant changes in tonic excitatory currents, synaptic transmission, resting membrane potential (RMP), and neuronal excitability. Furthermore, we demonstrated that knockout of GluD1 attenuates the effects of NASPM, highlighting the critical involvement of this receptor in regulating neuronal function in the dlBNST.

### Role of GluD1 in BNST

Our experiments provide evidence of a NASPM-sensitive tonic excitatory current in the mouse dlBNST (Figs. 1 and 4). The significant upward shift in dlBNST holding current following NASPM application in C57BL6/J and WT animals, but not in GluD1 KOs, suggests that GluD1 contributes to a baseline tonic excitatory current in these neurons. We found that NASPM induced hyperpolarization of the RMP (Fig. 3b-c), increased rheobase (Fig. 3d), and decreased the number of action potentials fired in response to escalating current steps (Fig. 3e), suggesting that tonic conductance via GluD1 plays an important role in regulating intrinsic neuronal excitability in the dlBNST. These data indicate that, like the dorsal raphe, GluD1 is constitutively active in the BNST and has functional ramifications on neuronal signaling (Gantz et al. 2020; Copeland et al. 2023).

GluD1 also contributes to synaptic excitatory signaling in the dlBNST. While sEPSC amplitude was unchanged, NASPM robustly reduced sEPSC frequency in all groups. While it remains possible that NASPM could be acting via presynaptic polyamine sensitive receptors, such as presynaptic NMDARs (Williams et al. 1991; Larsen et al. 2014), in the BNST we have shown no change in the paired-pulse ratio (PPR) following NASPM application (McElligott et al. 2010), suggesting that NASPM’s effects are postsynaptic.

This NASPM-induced decrease in sEPSC frequency was most pronounced in the GluD1 KO animals. Previous research has shown that GluD1 KO causes downregulation of GluA1 and GluA2 in the hippocampus and prefrontal cortex (PFC) (Yadav et al. 2012; Yadav et al. 2013). Loss of GluD1 also increases the frequency of mEPSCs in both regions (Gupta et al. 2015; Dai et al. 2021). In the PFC, loss of GluD1 coincides with an increase in the number of synapses and no change in amplitude (Gupta et al. 2015), while reports of synaptic density changes are more mixed in the hippocampus (Gupta et al. 2015; Dai et al. 2021). Conversely, GluD1 KOs exhibit upregulated GluA1 in the amygdala, while GluA2 expression is not affected (Yadav et al. 2012). Thus, the relationship between δGluRs and AMPARs is region- and likely cell type-specific. While we did not examine the effects of GluD1 KO on NMDARs in the BNST, it is likely that they are influencing the function of these receptors as well, and future experiments will explore these potential interactions.

### Role of GluD1 in Disease States

Variations in GluD1 have been associated with several neuropsychiatric and developmental disorders, including schizophrenia, bipolar disorder, autism spectrum disorder (Fallin et al. 2005; Griswold et al. 2012; Andrews and Dravid 2021), and substance use disorders (see below). In preclinical research, GluD1 KO mice display a number of behavioral abnormalities, including aberrant social interaction and hyperaggression, enhanced depressive-like behavior, reduced anxiety-like behavior, and deficits in learning and memory (Yadav et al. 2012; Yadav et al. 2013; Nakamoto et al. 2020a).

GluD1 has also been implicated in alcohol use disorder (AUD). One genome-wide association study identified GluD1 as linked to comorbid depression and AUD (Edwards et al. 2012), while another conducted in college students positively associated variations in GluD1 with the maximum number of drinks consumed in a 24 hour period across every racial background tested – the only gene to meet this criterion (Webb et al. 2017). Both GluD1 and GluD2 mRNA expression is enhanced in the caudate nucleus in postmortem brains of individuals with AUD while GluN2A is diminished, again demonstrating a relationship between δGluRs and synaptic ionotropic receptors (Bhandage et al. 2014). Additionally, GluD1 has been shown to be upregulated in mice bred for a high alcohol-drinking and low withdrawal-symptom severity phenotype compared to their low-drinking, high-withdrawal counterparts (Kozell et al. 2020). Deletion of striatal GluD1 in mice results in greater cocaine conditioned place preference (CPP) compared to WT controls, and leads to alterations in basal MSN NMDAR subunit composition and silent synapse formation, as well (Liu et al. 2018). Collectively, these studies suggest that δGluRs augment behavioral responses to reinforcing substances and identify δGluRs as a potential pharmacotherapeutic target for substance use disorders.

### Potential Role of BNST GluD1 in Disease States

The BNST is an intriguing nucleus that is anatomically positioned to integrate both appetitive and aversive sensory experiences, memories, and interoceptive states (Stamatakis et al. 2014; Lebow and Chen 2016). It plays a significant role in mediating addictive-like behaviors and is the only recognized node for anxiety disorders in the NIMH Research Domain Criteria (Koob 2008; Vranjkovic et al. 2017). The BNST also expresses mGluR1/5 and α1-AR, Gq-coupled receptors which have been shown to interact with GluD1 in other brain regions. Previous research from us and others has investigated glutamatergic signaling downstream from both of these receptors, including an α1-AR and an mGluR5-mediated LTD in the lateral BNST (McElligott and Winder 2008; Grueter et al. 2008; McElligott et al. 2010). Application of NASPM in slices from naive mice eliminated 30% of the evoked EPSC (eEPSC) response, but did not significantly affect EPSC amplitude in slices after α1-AR LTD had been pharmacologically induced, indicating that this α1-AR LTD is mediated by the removal of CP-AMPARs from the synapse (McElligott et al. 2010). Furthermore, this α1-AR LTD was attenuated following both chronic restraint stress and acute withdrawal from chronic intermittent ethanol vapor (CIE) exposure, and NASPM application reduction of eEPSC amplitude was attenuated in slices from chronic restraint stress-exposed mice (McElligott et al. 2010). These data indicate that CP-AMPARs are removed from the synapse following chronic stress *in vivo* via α1-AR LTD. Given that this α1-LTD is also blunted during CIE withdrawal, this suggests that withdrawal from alcohol similarly impacts the function of BNST CP-AMPARs. The impact of GluD1 in this plasticity, or if GluD1 function itself is impacted by these physiological challenges is currently unexplored and will be fruitful avenues for future research.

### Role of δGluRs in NASPM-Induced Effects

To our knowledge, this study and work from the Gantz lab are the only investigations to report the existence of tonic GluD1 conductance in the BNST and dorsal raphe, respectively. It is likely, however, that the change in holding current induced by NASPM in electrophysiological recordings is often missed. Indeed, our group has published several papers using NASPM as a pharmacological tool, but we previously overlooked changes in holding current during episodic stimulation (McElligott et al. 2010; Faccidomo et al. 2021; Faccidomo et al. 2024).

While NASPM is commonly regarded as a CP-AMPAR-selective antagonist, its ability to block δGluRs represents an extremely important consideration for the design of future studies, and augments the interpretation of previous results in publication, especially in cases of intracranial NASPM administration (Zhang et al. 2018; Faccidomo et al. 2021). For instance, Torquatto et al. observed that infusion of NASPM into the BLA or hippocampus produced impairments in fear memory consolidation and retrieval and deficits in reversal learning in the Morris water maze, while leaving object memory unchanged (Torquatto et al. 2019). As functional incorporation in GluA1 in BLA synapses plays an important role in fear conditioning (Rumpel et al. 2005), the investigators attributed these effects to NASPM suppression of CP-AMPAR activity; however, these deficits are phenocopied in GluD1 KO mice (Yadav et al. 2013). Certainly, changes to CP-AMPAR activation and synaptic trafficking are crucial components of neuronal adaptation. Yet given the role of δGluRs in regulating synaptic plasticity, as well as the open question of whether there is also G_q_PCR-facilitated or constitutive δGluR activity in yet unexplored brain regions, further examination of the potential role of δGluRs in these phenotypes may grant additional mechanistic insight.

In conclusion we have demonstrated here that GluD1 contributes to a tonic current in the dlBNST, and influences synaptic transmission and cell excitability of male mice. Future studies will unravel the role of this structural protein and functional ion channel in the BNST on behavior and downstream effector systems.

## Resources

Chemicals: Picrotoxin (25 μM PTX, Abcam), 1-naphthylacetyl spermine trihydrochloride (100 μM NASPM, Cayman Chemical), DMSO (Sigma), isoflurane Animals: C57/B6J (Jackson Laboratories), GluD1 KO and WT littermate controls bred from heterozygote x heterozygote crosses; starter mice were a gift from Dr. Stephanie Gantz (University of Iowa) and originally generated by Dr. Jian Zuo (Creighton University).

PCR Primers for KO validation: Transnetyx, PCR genotyping assay (30 cycles) with a pair of primers from the deleted region of the *grid1* gene (5′ GCAAGCGCTACATGGACTAC 3′ and 5′ GGCACTGTGCAGGGTGGCAG 3′) and a pair of primers from the targeting vector (5′ CCTGAATGAACTGCAGGACG 3′ and 5′ CGCTATGTCCTGATAGCGATC 3′) from ear tissue samples.

## Acknowledgements

We would like to thank Dr. Stephanie Gantz for the GluD1 KO starter mice and helpful conversations about δ glutamate receptors.

## Supplemental Figures

**Supplemental Table 1.** Membrane properties of dlBNST neurons in naïve male mice. Data are expressed as mean ± SEM.

| **Supplemental Table 1** Membrane properties of dLBNST neurons in naïve male mice. Data are expressed as mean ± SEM. | | | | |
| --- | --- | --- | --- | --- |
| | N | $R_m$ (MΩ) | $C_m$ (pF) | $R_a$ (MΩ) |
| C57BL6/J | 14 mice, 24 cells | 397.1 ± 43.16 | 48.18 ± 4.598 | 16.17 ± 0.9393 |
| GluD1 WT | 9 mice, 16 cells | 363.8 ± 52.35 | 65.66 ± 6.917 | 16.86 ± 0.9810 |
| GluD1 KO | 7 mice, 15 cells | 444.5 ± 57.92 | 52.21 ± 5.267 | 18.75 ± 1.188 |

**Supplemental Fig. 1.**
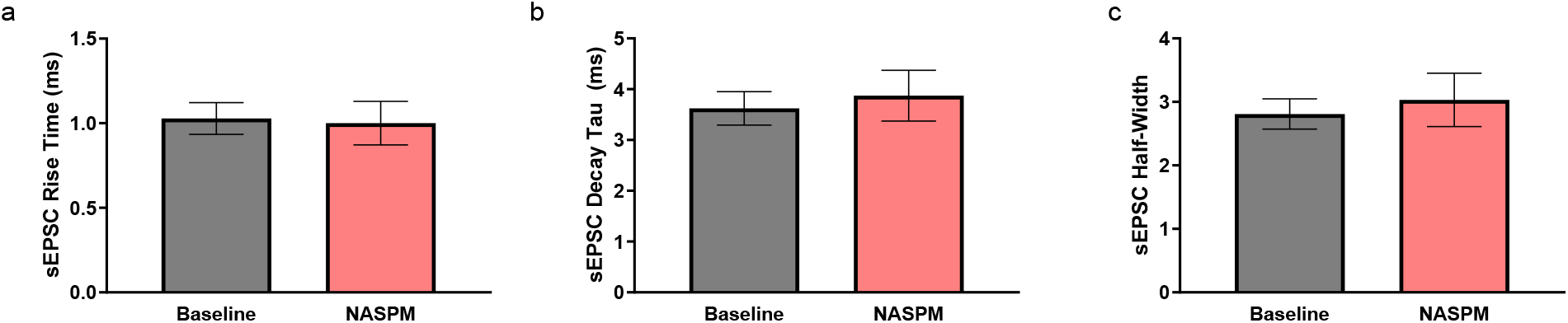
Naïve C57BL6/J sEPSC Kinetics. Paired t-tests did not show a significant change in sEPSC rise time (**a**), decay (**b**), or half-width (**c**) were observed before and after application of 100 µM NASPM in the dlBNST of male mice. Total n = 11 cells from 8 mice.

**Supplemental Fig. 2.**
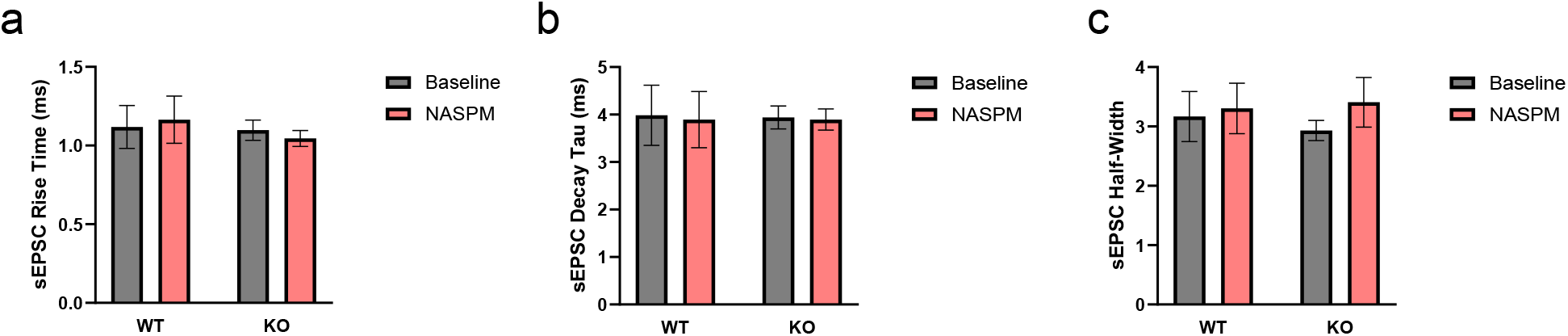
GluD1 WT and KO sEPSC Kinetics. No significant changes were observed in sEPSC rise time (**a**), decay (**b**), or half-width (**c**) in either genotype due to NASPM in a two-way RM ANOVA. Total n = 16 cells, 9 mice for WTs, 15 cells, 7 mice for GluD1 KOs.

